# Gene function prediction from bulk coexpression is bounded by cell-type-level signal

**DOI:** 10.64898/2026.08.31.748380

**Authors:** Alexander Adrian-Hamazaki, Paul Pavlidis

**Author notes:** Contributing authors.

## Abstract

It is widely accepted in genomics that coexpression of RNA transcripts suggests a commonality of function. This intuition is explicitly leveraged in machine learning methods that predict gene function, where it is often combined with other features such as protein interactions and sequence similarity. For example, including coexpression data from human tissue expression boosts performance for predicting Gene Ontology annotations. However, the biological underpinnings of this observation have not been well-investigated. Building on earlier results from our group, in this work we show that gene function is predictable from coexpression substantially because it reflects differences in expression between cell types, and these differences are also intrinsic to the ground truth labels. Using simulations and analyses of real data, we show that variance in the cellular composition of bulk samples impacts function learnability and attribute this to cell type marker gene content in the GO terms. We further show that cell type profiles, where the relationship between gene expression and cell type is made transparent, are effective for predicting gene function while increasing interpretability. These results indicate that function prediction models trained on bulk coexpression are largely limited to cell-type-level resolution rather than fine-grained biochemical function, with direct consequences for how such predictions should be interpreted.

## 1 Introduction

Predicting the functions of genes has long been a focus in bioinformatics. Originally, function prediction was performed using sequence similarity, with algorithms such as BLAST being widely adopted by the community [1]. With the advent of high-throughput methods, function prediction methods began using data modalities such as coexpression [2–5], protein-protein interaction [6] uni- and multi-modally [7–10], in the hopes of predicting gene functions through their “guilt-by-association” (GBA) instead of through primary structure. Unfortunately, despite decades of development, GBA models have not been widely adopted by biologists or function annotators. Performance improvements of these models appear stagnant, and models still struggle to predict functions that cannot be inferred from sequence [8, 11–13]. In previous work from our group, we identified multiple ways in which function prediction methods struggle to provide meaningful real-world performance, emphasizing problems with protein-protein interaction data [14]. Here we add to this body of work with an investigation of the contribution of coexpression data to mammalian gene function prediction. Although coexpression data showed early promise for function prediction and continues to be a commonly employed data modality, its underlying contribution to function prediction is poorly understood. In this paper, we aim to uncover principles that govern the predictive abilities and limitations of bulk coexpression.

Our particular focus is coexpression computed from bulk tissue samples, which has been relied on in most function prediction studies for genes of multicellular organisms, both as a standalone modality and as an input to integrative methods [7, 9, 10]. The logic of using coexpression to predict function is based on the assumption that genes expressed in a coordinated, co-varying fashion are more likely to be involved in shared cellular processes than non-coordinated genes (guilt-by-association) [2, 4]. The implicit hope is that a model trained on coexpression could uncover high-resolution biological gene function, such as involvement in protein complexes, metabolic pathways, or signal cascades occurring transiently or during experimental conditions (drug treatments, gene knockouts) [15–17]. The feasibility of this goal has been challenged at the level of network-wide guilt-by-association performance [13, 14, 18], but its biological underpinnings — why coexpression predicts function when it does — have not been directly investigated. Consequently, the extent to which models can glean high-resolution cellular information from bulk coexpression is unknown.

Coexpression modules derived from bulk tissue have long been observed to group cell type marker genes, so that modules correspond to major cell classes [19–21]. We have built on these observations, showing that cell type compositional differences across bulk samples induce covariance between the genes that have cell-type-specific expression patterns [22], which enhances co-expression among cell type-specific genes. Consequently, bulk coexpression is a combination of at least two sources of covariance: “composition-induced coexpression” and “cross-cell co-expression”. The former reflects differences in expression between cell types, whereas the latter reflects differences in expression between cells. Previously, we have estimated that up to 50% of bulk coexpression can be composition-induced in real data [22], and that in simulations, cross-cell co-expression becomes diluted by composition-induced co-expression [23]. Whether this effect represents a limitation on the information content of bulk coexpression to support high-resolution inference on function is unknown. If it does, it would be a major limitation, implying that such models capture coarse cell-type-level signal rather than high-resolution gene function.

Whether the cell type composition-induced signal helps or hinders function prediction depends critically on how gene function itself is defined — and in particular, on if the labels represent the information content in the composition-induced signal. The Gene Ontology (GO) annotations [24, 25] are the gold standard label set for function prediction and evaluation [8, 26–28]. We first observed that many GO terms, especially in the Biological Process ontology, have an element of low-resolution cell-type specificity (e.g. “GO:0006836, Neurotransmitter Transport”; “GO:0050852, T cell receptor signaling pathway”). We hypothesize that composition-induced coexpression signal in bulk coexpression strengthens predictions for GO terms that are cell-type-affiliated in name insofar as they are enriched for genes that express in cell-type specific patterns (Fig. 1). This could explain both why bulk coexpression performs poorly on most GO terms (they contain genes which are not cell-type specific), and why the usability for even highly predictable functions is still poor (the label is too cell-type specific and low-resolution).

**Fig. 1.**
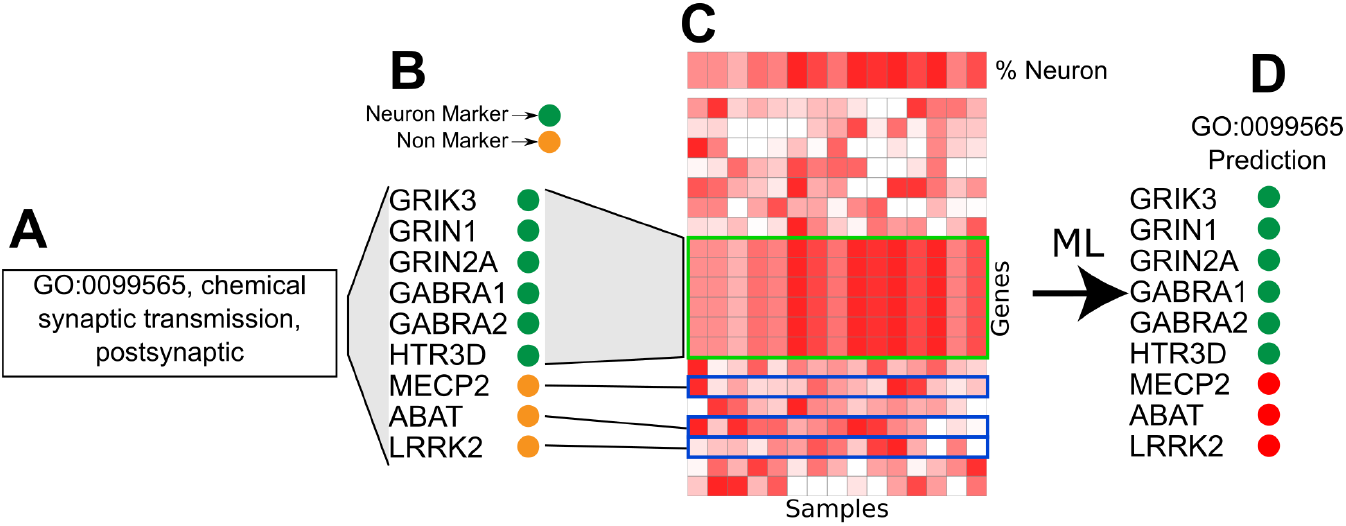
Hypothesized framework for how composition effect enables GO learnability. GO:0099565, a brain-related GO term (**A**) is annotated with neuron marker genes and non-marker genes (**B**). (**C**) Neuron composition variance across samples and its effect on coexpression among neuron marker genes (green box) and non-marker genes (blue boxes). (**D**) Guilt by association machine learning models trained on the resulting coexpression.

In this paper, we formally evaluate the extent to which composition-induced coexpression affects gene function prediction. Through a series of simulations and analyses of real data, we show that bulk coexpression is limited in part due to the presence of cell-type-driven covariance in the data, combined with the cell-type-level framing of many functions by GO. This suggests a lowering of expectations for function prediction from a high-resolution that yields insights into biochemical cellular events, to a cell-type level resolution. It also suggests that the phenomenon could be exploited to gain insight into cell-type-level resolution functions. We provide a demonstration of this idea using cell-type-specific expression profiles.

## 2 Methods

Analyses were conducted with custom Python and R scripts. Code to reproduce the analyses is available [here].

### 2.1 Data

#### 2.1.1 Bulk tissue RNA-seq data

Bulk RNAseq data was downloaded from GTEx [29] (GTEx analysis version 8). The dataset contains 17382 samples across 54 distinct tissue types. The dataset contains 2642 brain samples.

#### 2.1.2 Single cell RNA-seq data

Single-cell RNA-seq data was downloaded from the Human Protein Atlas [30] (accessed August 2023). A standard protocol was used to remove low-quality cells [31]. The dataset contains cells (Supplementary Table S1) from 31 tissues and a total of 85 cell types. It contains six major brain cell types (excitatory neurons, inhibitory neurons, microglial cells, astrocytes, oligodendrocytes, and oligodendrocyte precursor cells). For cell type profile analysis, we collapsed cells of the same cell type into a cell type profile by calculating each gene’s mean expression in the cell type. This yielded expression values for 19437 genes across 85 cell types.

#### 2.1.3 GO terms

GO biological gene annotations were downloaded from the GO (version 2.2) [24, 25]. Only terms with 20 to 200 genes were kept. When two GO terms shared greater than or equal to 70% of the same annotated genes, only one term was kept. This yielded 1396 GO terms (Supplementary Table S2).

### 2.2 Function prediction

All function prediction in this work was done using the EGAD algorithm as implemented in the EGAD R package [32]. EGAD is a simple, rapid, and high-performing prediction model that has been applied to coexpression analysis [32]. EGAD takes a gene network (in our case, a coexpression network), performs label propagation across 5 cross-validation folds, and evaluates how well the labels in the hidden fold are recapitulated via AUROC.

### 2.3 Simulating composition-induced coexpression

We established an algorithm to simulate bulk tissue RNAseq expression data whose only source of cross-sample variance is composition variance. Consequently, coexpression derived from these tissues is a pure measure of composition-induced coexpression. In the case of simulating brain composition-induced coexpression we follow the following steps:

1. We gathered the major type profiles for cells that comprise the brain cell types: excitatory and inhibitory neurons, astrocytes, microglial, oligodendrocytes, and oligodendrocyte precursors. Although in this work we studied composition-induced coexpression in brain, in principle this process could be followed for any tissue type.
2. For each cell type, we defined a biologically informed Gaussian distribution reflecting its plausible percent composition in bulk tissue (Eq. 1). These estimates (Supplementary Table S3) were histologically based [33, 34].

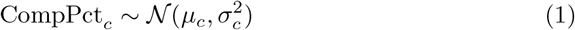

where CompPct_*c*_ is the percent composition of cell type *c, µ*_*c*_ is the mean percent composition of cell type *c*, and 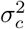 is the variance of the percent composition of cell type *c*.
3. To simulate one bulk brain sample, we sampled percent composition values for each cell type. The set of percentages was scaled to sum to one. Each cell type profile was then scaled according to its percent contribution to the sample. The resulting scaled profiles were summed, yielding the final bulk expression (Eq. 2).

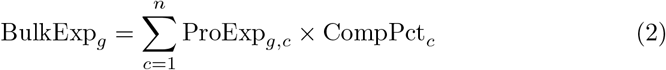

where BulkExp_*g*_ is the simulated bulk expression of gene *g*, ProExp_*g,c*_ is the expression level of gene *g* in cell type profile *c*, CompPct_*c*_ is the percent composition of cell type *c*, and *n* is the total number of cell types.

The general workflow is to simulate bulk expression using this algorithm, and then take coexpression from the simulated data to train function prediction models.

### 2.4 Simulated functions

To measure our hypothesis, we simulated cell-type-related functions that we would expect to be learnable from composition-induced coexpression. To do this, we first performed a differential expression analysis [31] on the single cell brain data. Then, for each cell type, we categorized these genes into three groups: top 5% of up-regulated genes, bottom 5% of up-regulated genes (down-regulated genes), and all other genes (non-associated). To simulate a function with genes enriched for a cell type — such as an excitatory neuron function — we sampled (with replacement) 20 upregulated genes. As a control, we also simulated non-associated functions that are comprised of 20 non-associated genes. These simulated functions served as ground truth labels for evaluating composition-induced learnability. The genes in the simulated terms can be found in Supplementary Table S4.

To simulate brain-related functions, we first calculated Gini indices of brain cell type profile expression across the six brain cell types. To simulate a function with brain cell-type related genes (brain-enriched), we sampled 20 genes from the top 5% of Gini indices. As controls we also simulated functions with genes not in the top 5% of Gini indices (non-brain enriched). The genes in the simulated terms can be found in Supplementary Table S5.

### 2.5 Marker gene content scores

We developed the Marker Gene Content (MGC) score as a way to quantify how tissue specific a GO term is. To calculate the MGC scores for a given tissue, we first identified which cell types pertain to that tissue. Next, we calculated the Gini indices of the gene expression profiles across those cell types. We define the MGC score for a GO term as the mean of the Gini indices of the genes comprising the term. In this work, we make use of brain-MGC scores, which were calculated using the cell type profiles of the aforementioned major brain cell type. For Pan-MGC scores, this process was performed for all cell type profiles.

### 2.6 Statistical comparisons

Several statistical comparisons were performed comparing model performance across different training datas. Standard two-tailed Student’s t tests were performed for some comparisons. Generalized linear mixed effects models (GLMMs) with Gaussian links were fit for analyses on the simulated GO terms, where the simulated functions were treated as random effects in the regression.

## 3 Results

### 3.1 Simulated composition-induced co-expression enables prediction

#### 3.1.1 Simulated GO terms are learnable

To show that composition-induced co-expression impacts function prediction in principle, we established a simulation framework for simulating pure composition-induced co-expression. Briefly, the method collapses single-cell RNAseq data into cell type profiles, thereby removing cross-cell-type variance, and samples the profiles, combining the profiles to create bulk RNAseq samples composed of different percentages of cell type profiles (Methods). We used this method to simulate bulk brain composition-induced coexpression datasets. We also created a simple method to create simulated functions whose gene membership is constrained by their level of expression in a given cell type (Methods). Using this method, we simulated a set of functions enriched for excitatory neuron genes, and a set containing non-excitatory neuron genes (Supplementary Table S4). Then, using EGAD [32], a neighbor-voting function prediction algorithm, we trained prediction models using the simulated brain composition-induced coexpression to predict gene membership of the simulated functions. We found that functions that were enriched with excitatory neuron-related genes were more learnable (Fig. 2a, median AUROC = 0.71) than those with non-associated genes (median AUROC = 0.50, generalized linear mixed model *p <* 2.16 *×* 10^*−*16^). To probe this relationship further, we investigated how removing the cell-type specific expression content from the training data impacted learnability. We simulated a set of bulk brain coexpression samples that lacked excitatory neuron expression content. This significantly deteriorated model performance on the simulated excitatory neuron functions (median AUROC = 0.53, generalized linear mixed model *p <* 2.16 *×* 10^*−*16^) (Fig. 2a). These results show that composition-induced coexpression provides predictive power for functions insofar as they are annotated with cell type-specific genes.

**Fig. 2.**
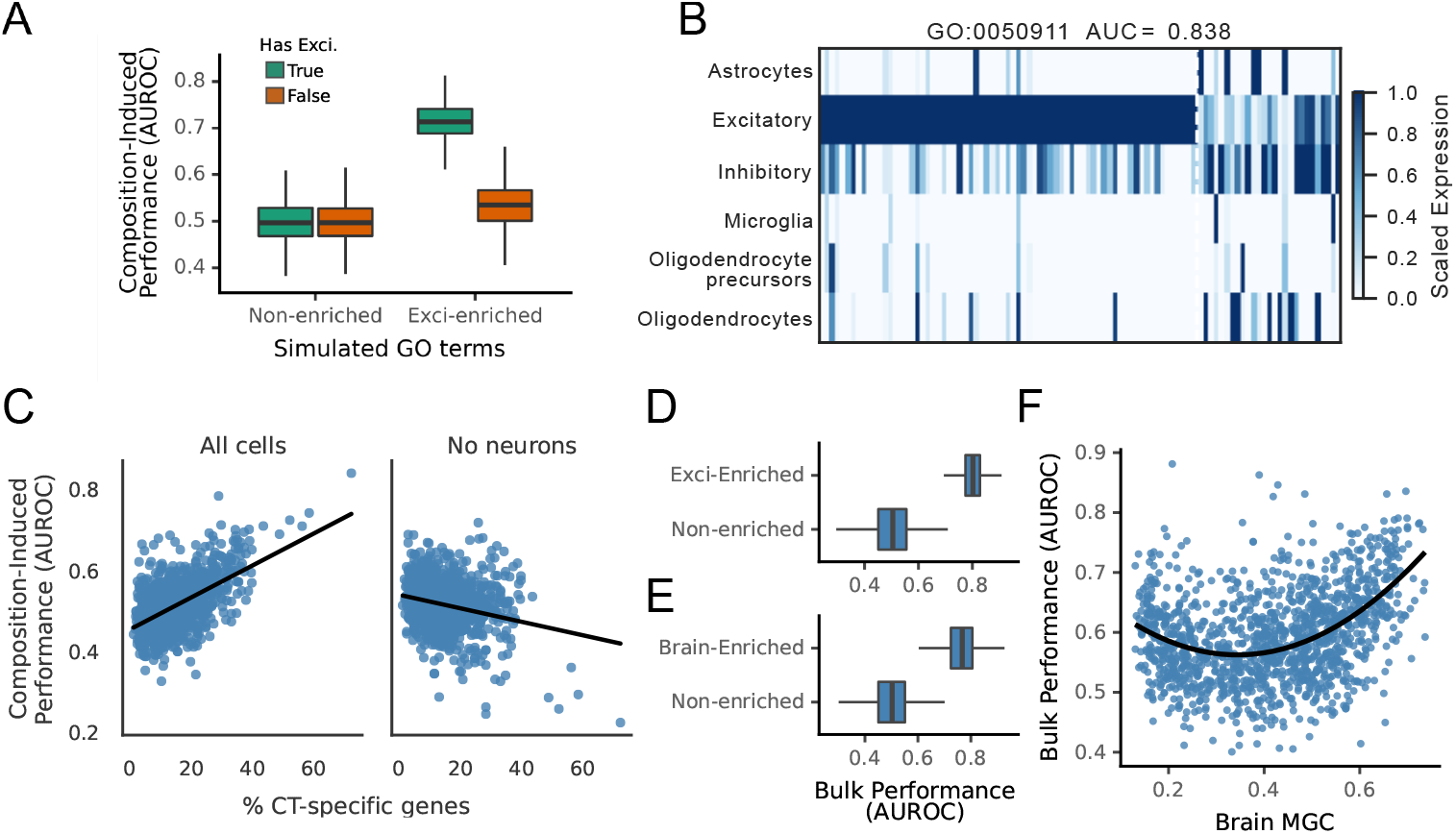
Composition-induced coexpression and cell-type-specific GO terms. (**A**) Prediction performance (AUROC) on simulated GO terms enriched for excitatory neuron genes (Exci-enriched) versus non-enriched terms, when training data either includes or excludes excitatory neuron expression content. (**B**) Expression heatmap of the 98 genes annotated in GO:00050911 across six major brain cell types. (**C**) Relationship between GO term learnability and the percentage of cell-type-specific genes in the term under two training conditions – all cell types included (left) or excluding neurons (right). (**D**) Learnability of simulated excitatory-neuron-enriched versus non-enriched GO terms when models are trained on real bulk brain coexpression. (**E**) Learnability of simulated brain-enriched versus non-enriched GO terms under the same training data. (**F**) Relationship between real GO term learnability and Brain Marker Gene Content (MGC) score for models trained on bulk brain coexpression, with a LOESS smoothing curve overlaid.

#### 3.1.2 Real GO terms are learnable from composition signals

Next, we investigated the extent to which this effect is present for real GO terms trained on simulated data. We trained models on simulated composition-induced coexpression to predict gene content of 1396 high-confidence (Methods) Biological Process GO terms. We first observed that many of the top performing GO terms contained genes highly expressed in excitatory and inhibitory neurons. For example, GO:00050911 (Fig. 2b), the highest performing term (AUROC = 0.84) was annotated with 98 genes, 71 of which were highly expressed in excitatory neurons relative to the other cell types; inhibitory neuron expression in this GO term was highly correlated with excitatory neuron expression (Pearson *R*^2^ = 0.94), thus they were combined in subsequent analysis. When we trained models on composition-induced coexpression that lacked neuron content, GO:00050911 was no longer learnable (AUROC = 0.22). To formalize this observation, we asked if learnability could be explained by how much a GO term is dominated with genes highly expressed in one cell type. To measure this, for each gene, we annotated which cell type had the highest expression of that gene, and we used these annotations to identify the fraction of genes in each GO term that pertained to each cell type. Then, for each cell type, we regressed GO term learnability against the fraction of genes in the term that pertained to that cell type. We found that GO terms with a large fraction of neuron-related genes tended to be more learnable when models were provided neuron expression information (0.03 AUROC/10% increase, Fig. 2c), however, this relationship was attenuated when the models were not provided neuron expression information (*−*0.01 AUROC/10% increase).

Having shown that removing neural information from training decreases performance of neuron-related GO terms, we asked how removing neural information impacts non-neural GO terms. In non-neuronal cell types, when models were trained with all cell types, we did not observe a strong positive relationship between learnability and cell-type-specific gene content (Fig. S1). We speculated that this was due to a general lack of cell type specific gene content in the functions (Fig. S1). Microglial, astrocytes, and oligodendrocytes, had negative or neutral relationships (*−*0.02, *−*0.006,*−*0.01 AUROC/10% increase respectively) (Fig. S1). However, when excitatory neurons were removed from the training data, the relationships for these cell types all increased (0.003, 0.01, 0.02 AUROC/10% increase respectively), indicating that excitatory neuron genes — which are overrepresented in the expression data — may mask the relationship for other cell types. In support of our hypothesis, overall, these results demonstrate that real GO term learnability is enhanced by composition-induced coexpression when the GO terms are enriched for cell type specific genes.

### 3.2 Composition-induced co-expression explains performance in bulk co-expression

Based on our simulations, we speculated that the natural composition-induced coexpression in real bulk coexpression would contribute to prediction performance for GO terms enriched for cell type-related genes. We first probed this relationship by training models using real bulk brain coexpression data to predict gene membership of the simulated excitatory-neuron-related GO terms. The terms enriched for excitatory neuron genes were more learnable (mean AUROC = 0.80), compared to the terms that contained non-excitatory neuronal genes (mean AUROC = 0.50, Student’s *t p <* 2.16 *×* 10^*−*16^, Fig. 2d). However, given that bulk data contains composition-induced coexpression, and that we expect the compositional differences to exacerbate covariance for *all* cell types in the bulk, we hypothesized that GO terms that contained cell type-related genes for *any* brain cell type (in other words, GO terms that are brain-related) would be more learnable. To test this, we developed the Marker Gene Content (MGC) score, which quantifies a GO term’s brain-relatedness by averaging the Gini indices of its constituent genes’ expression across cell types (Methods), where a high score indicates GO terms with more genes whose expression highly varies across cell types. Following our method of simulated excitatory neuron related GO terms, we used the MGC to simulate brain-related GO terms, and non-brain-related GO terms. We found that models trained with bulk brain coexpression were significantly better at brain-related (high-MGC scores) functions (mean AUROC = 0.76) compared to the null (mean AUROC = 0.50, Student’s *t p <* 2.16 *×* 10^*−*16^) (Fig. 2e).

Having established that bulk coexpression increases learnability for brain-related GO terms in simulation, we asked whether the relationship would hold for real GO terms. We trained models on real bulk brain coexpression and compared each GO term’s learnability against its brain MGC score, finding that terms with high MGC scores were indeed more learnable (Fig. 2f). Taken together, these results suggest that real GO terms are learned in part because of composition-variance in bulk coexpression.

### 3.3 Cell type profiles recapitulate bulk performance

Having shown that performance on cell type-related functions depends on the presence of cell type-specific expression, we investigated whether coexpression derived directly from cell type profiles could be used to predict function while making the source of the biological signal explicit. Consistent with composition-induced coexpression, brain-related terms enriched for brain genes achieved significantly higher performance (Fig. 3a; mean AUROC = 0.68) than non-enriched terms (mean AUROC = 0.50; Student’s *t*-test *p <* 2.16 *×* 10^*−*16^). Performance was also highly concordant with models trained on composition-induced coexpression regardless of enrichment status (Fig. 3b; brain-enriched *R*^2^ = 0.72, non-enriched *R*^2^ = 0.71; both *p <* 2.16 *×* 10^*−*16^). Similarly, performance across real GO terms was strongly correlated between the two training paradigms (Fig. 3c; *R*^2^ = 0.83, *p <* 2.16 *×* 10^*−*16^). Together, these results indicate that cell type profiles capture much of the same functional signal present in composition-induced coexpression.

**Fig. 3.**
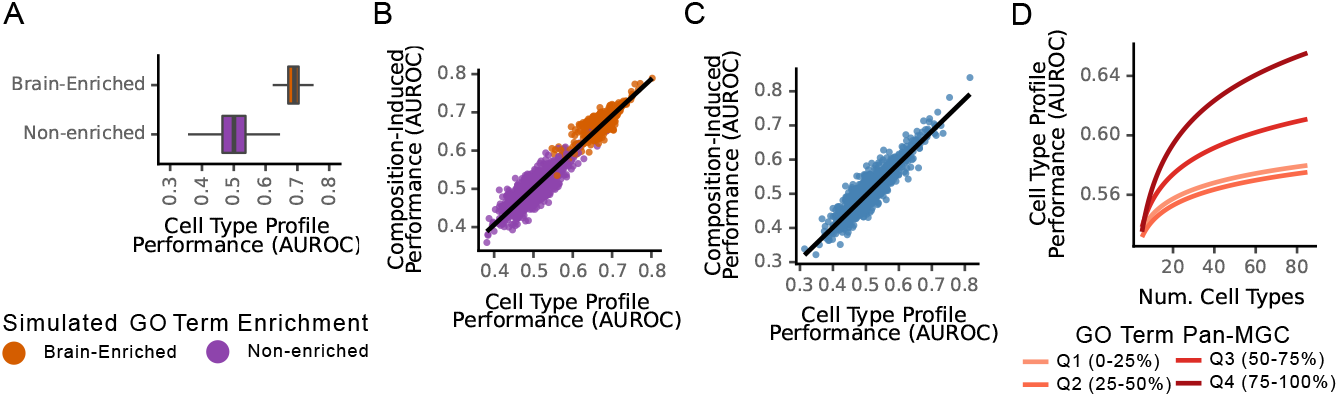
Cell type profiles and the predictive signal of composition-induced coexpression. Learnability (AUROC) of brain-enriched versus non-enriched simulated GO terms when models are trained on coexpression derived from only six brain cell type profiles. (**B**) Correlation between GO term learnability under composition-induced coexpression training versus cell type profile training, stratified by enrichment status (brain-enriched in orange, non-enriched in purple). (**C**) Correlation between GO term learnability under composition-induced coexpression versus cell type profile training across all real GO Biological Process terms. (**D**) GO term learnability as a function of the number of cell type profiles included in training, stratified by Pan-MGC score quartile (Q1: lowest 25%, Q4: highest 25%).

To further evaluate the extent to which cell type structure contributes to learnability, we investigated how varying the amount of cell type information available during training influenced model performance. We systematically increased the number of cell type profiles used in training to learn the real GO terms, and observed improvements in learnability as additional cell type information was incorporated (Fig. 3d); the extent of improvement was modulated by the GO term’s Pan-MGC score (Methods), suggesting that functions enriched for cell type specific genes benefited most strongly from the inclusion of additional cell type profiles. Finally, to assess how much signal in bulk coexpression could be explained by cell type structure alone, we compared models trained using coexpression from 85 cell type profiles against models trained on bulk coexpression from 54 tissue types. Learnability between the two approaches was strongly correlated (Pearson *R*^2^ = 0.67, *p <* 2.16 *×* 10^*−*16^), indicating that a substantial component of bulk coexpression performance may arise from underlying cell type-specific expression patterns. Together, these results suggest that merely informing models which genes are cell-type specific can explain predictive signal captured by bulk coexpression.

## 4 Discussion

Our goal in this work was to illuminate a key limitation in computational gene function prediction from bulk coexpression data. Using simulations and analysis of real data, we show that both the amount of cell-type specific content in the expression data *and* in the function are key modulators of model performance. This effect is a direct consequence of composition-induced coexpression in the bulk data, which enhances variance across the same cell-type specific genes present in the GO terms. Consequently, learnable functions tend to be those annotated with higher proportions of genes unique to one cell type. These results suggest a lowering of expectations for function prediction: from the ability to predict high-resolution intracellular functions, such as a gene’s involvement in a signalling pathway or protein complex, to a lower-resolution characterization of a gene’s activity in a specific cell type. These low-resolution predictions merely reflect cell-type specific expression differences rather than complex biochemical biology, limiting their utility. Our results indicate that non-cell-type-specific functions will be difficult to predict from bulk coexpression, and these observations should guide function prediction method developers and users to carefully consider how they interpret their results.

We stress that we are not claiming that cell-type-specific expression is never informative about gene function. Rather, we argue that if one is interested in determining the function of a gene that is highly expressed in a particular cell type, such as a gene expressed in excitatory neurons, it is already apparent that the gene may be involved in a neuron-specific function. A function prediction method is not needed to determine this. Still, our results hint that coexpression measured across cells of the same cell type could be leveraged to decouple composition-induced coexpression from intracellular coexpression and enable high-resolution predictions within that cell type. Such models would be beneficial for biologists insofar as predicting gene membership in high-resolution biochemical pathways and complexes is useful. Lending credence to this idea, recent work in our lab has shown that coexpression from single-cell data from a single cell type, both in theory and in practice, can more faithfully capture regulatory interactions than bulk tissue [23]. Lending credence to this, we note that some limited experimentally validated high-resolution information has been gleaned from guilt-by-assosiation in unicellular organism such as yeast [35, 36]. However, the extent to which this could be done with mammalian cells remains unclear, as the resolution of the predictions is fundamentally limited by the resolution of the labels.

These considerations bear directly on a body of work that uses tissue-specific functional networks to make gene-level disease inferences. Networks of this kind, built by integrating coexpression with other genomic data across human tissues, have been used to identify the changing functional roles of genes across tissues and to relate them to disease [37–42]. For example, a brain-specific network has been used to predict autism risk genes genome-wide [38]. Our results do not contest the observation that such networks rank neuronally expressed genes highly. They suggest a different reading of what that ranking represents. Because learnability from bulk brain coexpression tracks a term’s marker gene content (Fig. 2f), a candidate set derived from a brain network will be enriched for genes with cell-type-specific expression, and the annotations such a set converges on will preferentially be those that are themselves cell-type-affiliated. Convergence of this kind partly restates which cell types the input genes mark, rather than supplying independent evidence of a shared mechanism.

The main limitation of this work is our focus on brain tissue, without consideration of other tissue types. It is theoretically possible that other tissues would behave differently; however, composition variance is a factor in any bulk-sample-based profiling data, and we therefore expect the general principle to hold broadly. This expectation is supported by the fact that models trained on all 85 cell type profiles perform similarly to models trained on cross-tissue bulk data. A second limitation is our use of GO as the sole functional annotation scheme, without consideration of alternatives such as KEGG. However, to our knowledge, all such schemes contain functions with varying amounts of cell-type specific gene content, and our concerns would therefore likely apply to them as well. Finally, we used a single prediction method, EGAD, which we selected because it is rapid, high-performing, and relatively simple, with no free parameters to tune, reducing the risk of overfitting. Given its performance, and that all comparisons in this work use the same method, we do not expect this choice to alter our conclusions.

**Supplementary Fig. S1.**
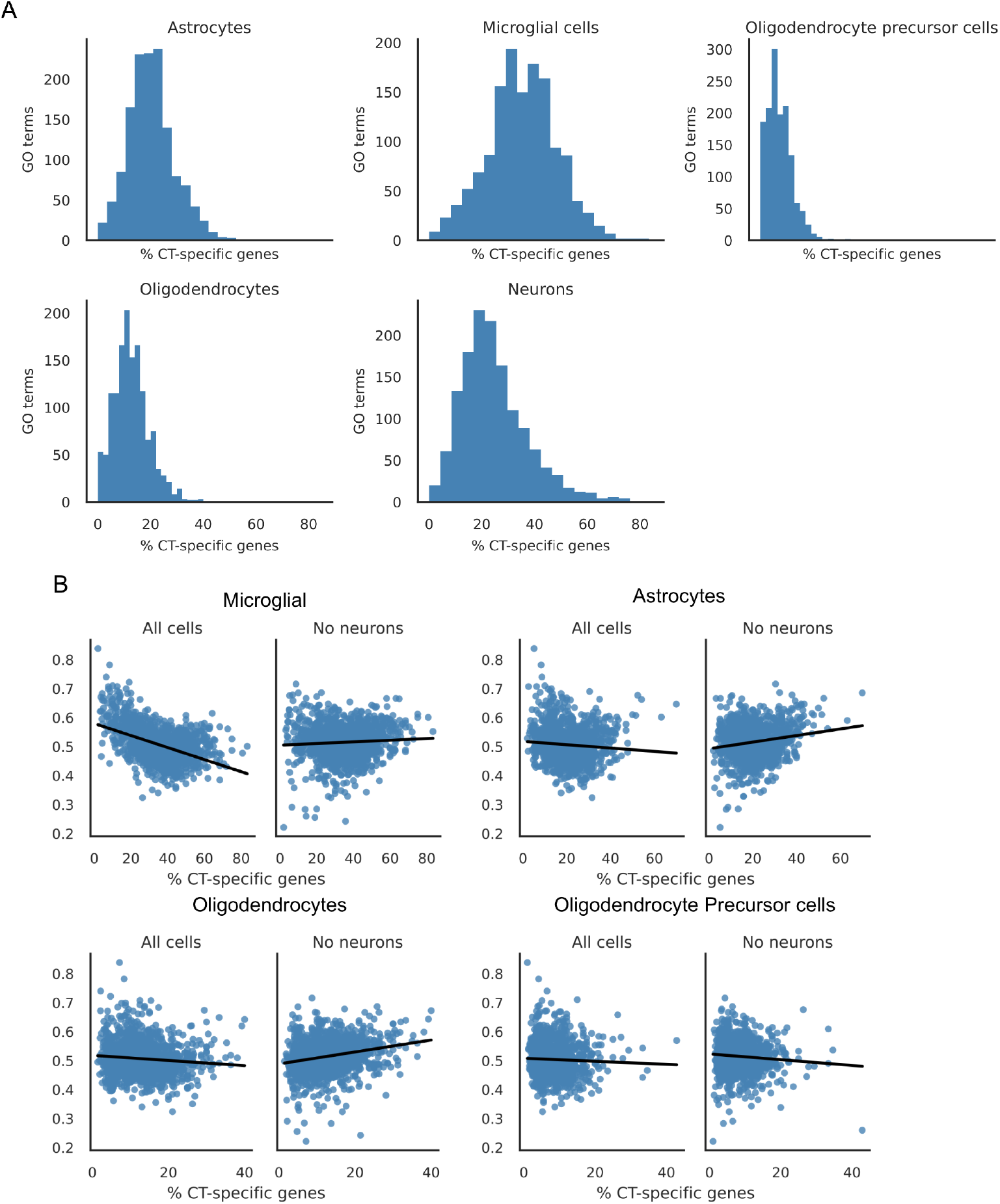
Cell-type-specific gene content across GO terms and its relationship to learnability for non-neuronal cell types. (**A**) Histograms of the percentage of cell-type-specific genes per GO term for each major brain cell type. (**B**) Relationship between GO term learnability and percentage of cell-type-specific genes for microglial cells, astrocytes, oligodendrocytes, and oligodendrocyte precursor cells, under two training conditions: all cell types included or excitatory neurons excluded.

## Acknowledgements

The authors thank Qinkai Wu, a previous student in the lab, for conducting preliminary experiments for this work. Funding sources are listed under Declarations.

## Declarations

### Ethics approval and consent to participate

Not applicable. This study used only publicly available, de-identified datasets.

### Consent for publication

Not applicable.

### Data availability

This study used publicly available data: bulk RNA-seq from GTEx (analysis version 8) and single-cell RNA-seq from the Human Protein Atlas. Gene Ontology annotations were obtained from the Gene Ontology Consortium (version 2.2).

### Materials availability

Not applicable.

### Code availability

Code to reproduce the analyses is available [here].

### Author contribution

A.A.-H. contributed to conceptualization, methodology, formal analysis, and writing. P.P. contributed to conceptualization, supervision, and review and editing. Both authors read and approved the final manuscript.

